# Aromatic Microbial Metabolite Hippuric Acid Potentiates Pro-Inflammatory Responses in Macrophages through TLR-MyD88 Signaling and Lipid Remodeling

**DOI:** 10.1101/2025.05.19.654724

**Authors:** Gauri Mirji, Sajad Ahmad Bhat, Mohamed El Sayed, Sarah Kim Reiser, Siva Pushpa Gavara, Ying Ye, Taito Miyamoto, Qin Liu, Aaron R Goldman, Andrew Kossenkov, Nan Zhang, Rahul S Shinde

**Author notes:** Correspondence to: Dr. Rahul S. Shinde, Molecular and Cellular Oncogenesis Program, Ellen and Ronald Caplan Cancer Center, The Wistar Institute, 3601 Spruce Street, Room 232, Philadelphia, PA 19104, USA.

## Abstract

The gut microbiome generates a diverse array of metabolites that actively shape host immunity, yet the pro-inflammatory potential of microbial metabolites remains poorly understood. In this study, we identified hippuric acid, an aromatic gut microbe-derived metabolite, as a potent enhancer of pro-inflammatory responses using a murine bacterial infection model and a non-targeted liquid chromatography-tandem mass spectrometry (LC-MS/MS)-based metabolomics. Administering hippuric acid intraperitoneally in murine models of *Escherichia coli* infection or LPS-induced inflammation significantly heightened pro-inflammatory responses and innate immune cell activation. *In vitro*, hippuric acid selectively potentiated M1-like macrophage polarization (LPS + IFNγ) but had no effect on M2-like polarization (IL-4). Hippuric acid further enhanced responses to diverse MyD88-dependent TLR ligands, but not TRIF-dependent TLR3, implicating a possible mechanism of action via activation of TLR-MyD88 signaling. Genetic deletion of MyD88 abrogated the pro-inflammatory effects of hippuric acid both *in vitro* and *in vivo*, confirming its dependence on the MyD88 pathway. Transcriptomic and lipidomic analyses revealed that hippuric acid promoted cholesterol biosynthesis and lipid accumulation, linking microbial metabolism to lipid-driven immune activation. Notably, hippuric acid similarly enhanced pro-inflammatory responses in human macrophages, and its elevated levels correlated with increased sepsis mortality, highlighting its potential clinical relevance. These findings establish hippuric acid as a previously unrecognized microbial-derived inflammatory modulator, bridging gut microbial metabolism, lipid remodeling, and innate immune signaling, and offer new insights into its role in infection and inflammation.

## Introduction

The gut microbiome is a vast and dynamic ecosystem that profoundly influences host physiology, particularly innate immune responses. A growing body of evidence highlights gut microbiota-derived metabolites as critical modulators of immunity, fine-tuning immune cell function and shaping both protective and pathogenic immune outcomes ^1,2^. These metabolites can enter systemic circulation and directly interact with immune cells in distant tissues, exerting broad immunomodulatory effects. While short-chain fatty acids (SCFAs), bile acid derivatives, and tryptophan metabolites have been widely characterized for their anti-inflammatory properties, other microbial metabolites such as trimethylamine N-oxide (TMAO) and p-cresol sulfate have been implicated in pro-inflammatory pathways ^3–9^. Despite these insights, the extent to which microbial metabolites actively enhance inflammation, particularly in the context of infection and sterile inflammation, remains poorly understood.

Among the diverse metabolic byproducts of the gut microbiota, aromatic metabolites derived from dietary and microbial metabolism have emerged as biomarkers and potential modulators of immune-related disorders. Hippuric acid, an aromatic microbial metabolite, has been associated with metabolic and inflammatory disorders, including obesity, cancer, and kidney dysfunction ^10–13^. It is synthesized through a two-step process: gut bacteria degrade dietary polyphenols (found in blueberries, tea, and other plant-based foods) into benzoic acid, which is subsequently conjugated with glycine in the liver to form hippuric acid ^14,15^. Despite its presence in circulation and its reported associations with immune-related diseases, the immunological function of hippuric acid has remained largely unexplored. Whether hippuric acid actively contributes to immune activation and inflammation has never been systematically investigated, particularly in the context of host-pathogen interactions and macrophage-driven immunity.

As central effectors of innate immunity, macrophages exhibit remarkable plasticity in response to environmental cues, polarizing into either pro-inflammatory (M1-like) or anti-inflammatory (M2-like) states ^16,17^. Traditionally, macrophages activated by lipopolysaccharide (LPS) and interferon-gamma (IFNγ) *in vitro*, termed as M1-like macrophages, secrete pro-inflammatory cytokines such as IL-12, TNFα, and IL-6, as well as generate reactive oxygen species (ROS) and nitric oxide (NO) to combat infections. In contrast, macrophages activated by IL-4 and IL-13 *in vitro*, termed as M2-like-like macrophages, promote tissue repair and immune resolution by upregulating markers like Arginase 1 and CD206. A tightly regulated balance between these macrophage phenotypes is crucial for immune homeostasis, and dysregulated macrophage activation contributes to inflammatory diseases. While microbial metabolites are increasingly recognized as modulators of macrophage polarization, their identity and mechanisms of action remain largely unknown, particularly in infection-induced inflammation.

Toll-like receptor (TLR)-MyD88 signaling is a central pathway linking microbial recognition to macrophage activation. TLRs recognize pathogen-associated molecular patterns (PAMPs) and endogenous danger signals, triggering MyD88-dependent activation of NFκB and MAPK pathways, which leads to the production of pro-inflammatory cytokines ^18–21^. Although TLR activation is classically initiated by microbial ligands such as LPS and bacterial lipoproteins, recent studies suggest that endogenous and microbial-derived metabolites may act as immunomodulatory signals that amplify or constrain TLR signaling. Furthermore, TLR activation is intimately connected to lipid metabolism, as inflammatory responses require substantial membrane remodeling and cholesterol biosynthesis ^22–24^. Accumulation of lipids can sustain macrophage activation and drive chronic inflammation, contributing to diseases such as atherosclerosis, autoimmunity, and sepsis ^25^. Metabolic rewiring of macrophages, particularly the changes in cholesterol biosynthesis, directly influences TLR signaling and cytokine production ^26^. However, which gut microbial metabolites intersect, and to what extent, with the TLR-MyD88 signaling pathway and drive macrophage lipid remodeling to potentiate inflammation remains unknown. Identifying pro-inflammatory microbial metabolites could fundamentally reshape our understanding of infection-induced immune dysregulation and chronic inflammatory diseases. Moreover, elucidating their mechanisms of action may uncover new therapeutic targets, allowing for metabolic interventions to modulate immune responses in a disease-specific manner.

Here, we identified hippuric acid as a potent enhancer of pro-inflammatory macrophage responses through TLR-MyD88 signaling and lipid remodeling. Using non-targeted metabolomics and inflammatory models, we demonstrated that hippuric acid selectively amplified M1-like macrophage polarization, enhanced inflammatory responses to diverse MyD88-dependent TLR ligands, and promoted cholesterol biosynthesis, thereby sustaining macrophage activation. Importantly, genetic deletion of MyD88 abrogated these effects, confirming the role of this pathway in hippuric acid-mediated inflammation. Furthermore, elevated hippuric acid levels correlated with increased mortality in sepsis patients, highlighting its clinical relevance in inflammatory diseases. These findings uncover a previously unrecognized role for microbial-derived hippuric acid in macrophage-driven inflammation and illuminate the intricate crosstalk between microbial metabolism, lipid signaling, and innate immune activation.

## Results

### Hippuric acid levels decreased during bacterial infection

The gut microbiome influences host immunity through microbial metabolites that modulate innate immune responses, macrophage polarization, and pro-inflammatory cytokine production. Bacterial infection serves as a robust model to study these interactions by inducing strong innate immune activation through pathogen-associated molecular patterns (PAMPs) that drive Toll-like receptor (TLR) signaling and cytokine production, such as IL-12p40 and IL-6 ^18^. To investigate changes in microbial metabolites during bacterial infection, C57BL/6 mice were infected intraperitoneally with *Escherichia coli* (2 × 10⁷ CFU), and non-targeted metabolomics using liquid chromatography-tandem mass spectrometry (LC-MS/MS) was performed on sera collected 48 hours post-infection (Figure 1A). The metabolomics screen revealed significant shifts in metabolic profiles, including altered levels of 2/3-hydroxybutyrate (a marker of fatty acid metabolism), hypoxanthine (a marker of purine metabolism), and N-acetylglutamine (involved in nitrogen metabolism), indicating broad metabolic reprogramming during infection (Figure 1B). Notably, hippuric acid, an aromatic microbial metabolite, emerged as one of the most significantly decreased microbial metabolites, with an average reduction of more than 24-fold in infected mice compared to controls (p < 0.001, Figure 1B-C). This finding suggests a strong link between bacterial infection and the depletion of hippuric acid, raising the question of whether this metabolite plays a role in modulating immune responses during infection.

**Figure 1:**
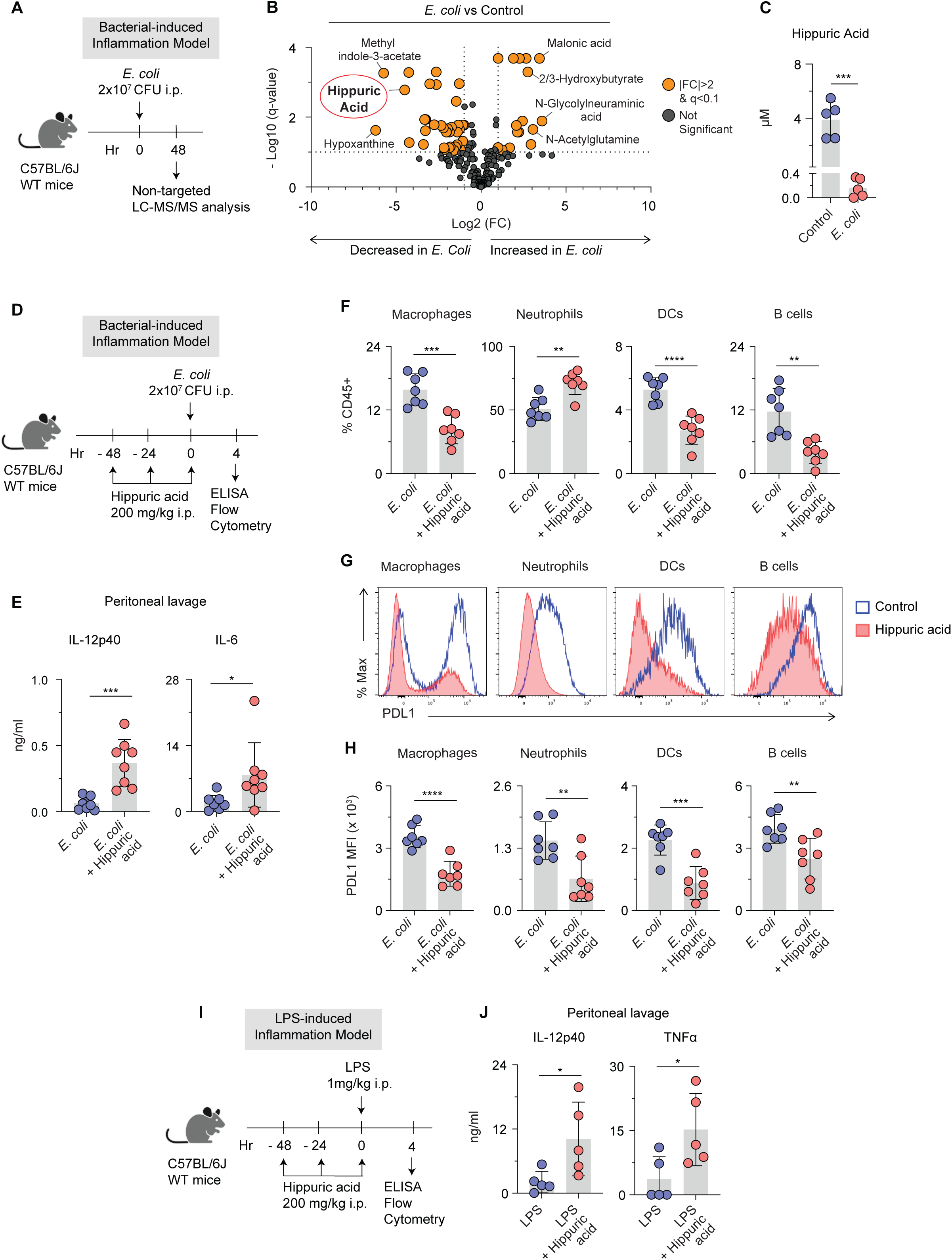
Hippuric acid potentiates pro-inflammatory responses in murine models of *E. coli* infection and LPS-induced inflammation. (A) Schematic of *E. coli* infection model for non-targeted metabolomics. (B) Volcano plot comparing metabolite profiles of sera from *E. coli* infected C57BL/6 mice 48 hr after *E. coli* inoculation in the peritoneum. Metabolomic analysis was performed using non-targeted liquid chromatography-tandem mass spectrometry (LC-MS/MS). n=5 mice per group. (C) Absolute quantification of hippuric acid in sera from *E. coli* infected C57BL/6 mice as in (A). n=5 mice per group. (D) Schematic of *E. coli* infection model showing time intervals for administering hippuric acid intraperitoneally and readouts of the study. (E) ELISA on peritoneal lavage obtained from mice in (D) for measurements of pro-inflammatory cytokines IL-12p40 and IL-6. n=8 mice per group. (F-H) Flow cytometry analyses of CD45+ immune cells in the peritoneum obtained from mice in (D) for changes in indicated immune cells as assessed for (F) percent populations, (G) histograms showing mean fluorescent intensity (MFI) of PDL1, (H) expression levels of PDL1 as MFI. n=7 mice per group. (I) Schematic of LPS-induced inflammation model showing time intervals for administering hippuric acid intraperitoneally and readouts of the study. (J) ELISA on peritoneal lavage obtained from mice in (I) for measurements of pro-inflammatory cytokines IL-12p40 and TNFα. n=6 mice per group. In (B), significant change was defined as |FC| > 2; *q* < 0.1 (Benjamini-Hochberg FDR-adjusted *P* value). In (C), (E-F), (H), and (J), data are presented as means ± SD and *P* values were determined by two-tailed Student’s *t-tests*.

### Hippuric acid potentiated pro-inflammatory responses during *E. coli* infection

To determine whether hippuric acid influences immune responses during bacterial infection, we administered hippuric acid intraperitoneally 48 hours prior to *E. coli* infection and assessed inflammatory responses 4 hours post-infection (Figure 1D). ELISA of peritoneal lavage fluid revealed a significant increase in levels of pro-inflammatory cytokines such as IL-12p40 and IL-6 in mice infected with *E. coli* receiving hippuric acid-treatment compared to those receiving *E. coli* infection alone (Figure 1E). Flow cytometry of CD45⁺ immune cells from the peritoneal lavage fluid showed that hippuric acid administration altered the immune cell populations, with a decrease in the proportion of macrophages, dendritic cells (DCs), and B cells but a marked increase in neutrophils (Figure 1F). Furthermore, their activation status shifted toward a pro-inflammatory phenotype. PDL1 expression, an immune checkpoint molecule associated with anti-inflammatory responses, was significantly reduced on macrophages, DCs, neutrophils, and B cells (Figure 1G-H). Additionally, there was a marked decrease in the expression of anti-inflammatory markers Arg1 (on macrophages) and CD206 (on neutrophils) (Supplementary Figure 1A-B), alongside an increase in expression of pro-inflammatory markers such as IL-6 on macrophages, CD86 on neutrophils, IL1β and TNFα on DCs, and CD86 and CD40 on B cells (Supplementary Figure 1A-D).

Given that bacterial infection often associates with systemic changes in immune responses, we next assessed phenotypic changes in immune cells from peripheral organs and lymphoid tissues (bone marrow and spleen). Flow analysis found a few differences in frequencies or phenotype of immune cells. Specifically, macrophage populations increased in the bone marrow, while neutrophils and DCs decreased in the spleen (Supplementary Figure 1E-H). PDL1 expression was similarly reduced on macrophages in the spleen, indicating that hippuric acid may influence systemic immune tolerance mechanisms (Supplementary Figure 1F-H). Taken together, these results suggest that hippuric acid amplifies pro-inflammatory cytokine production and immune cell activation during bacterial infection, while reducing markers of immune exhaustion, thereby acting as a pro-inflammatory metabolite.

### Hippuric acid amplified LPS-Induced inflammation

To determine whether hippuric acid broadly potentiates pro-inflammatory responses beyond bacterial infection, we utilized an LPS-induced inflammation model. Sub-lethal doses of LPS (1 mg/kg intraperitoneally) were administered following intraperitoneal pre-treatment with hippuric acid (Figure 1I). ELISA of peritoneal lavage fluid revealed a significant elevation in pro-inflammatory cytokines IL-12p40 and TNFα in mice treated with hippuric acid compared to LPS alone (Figure 1J). Flow cytometry of CD45⁺ immune cells demonstrated that hippuric acid enhanced the expression of immunostimulatory markers, including CD86 and IL-6 on macrophages, IL1β on neutrophils, and TNFα and IL-12p40 on DCs, while suppressing the production of the anti-inflammatory cytokine IL-10 by neutrophils cells (Supplementary Figure 1I-K). These findings suggest that hippuric acid primes the immune system to amplify TLR-mediated inflammation, further supporting its role as a metabolite that drives pro-inflammatory cytokine production and innate immune cell activation.

### Hippuric acid promoted M1-like macrophage polarization but showed no effect on M2-like polarization

We next investigated the direct effects of hippuric acid on macrophage pro-inflammatory responses. To test this, we used an *in vitro* model of M1-like polarization by stimulating bone marrow-derived macrophages (BMDMs) with LPS or LPS+IFNγ, both of which elicit strong pro-inflammatory responses. BMDMs were pre-treated with hippuric acid (10 µM or 100 µM) for 1 hour, followed by stimulation with LPS or LPS+IFNγ for 8 hours. ELISA of culture supernatants revealed that hippuric acid significantly increased the production of the pro-inflammatory cytokines IL-12p40 and IL-6 compared to either LPS or LPS+IFNγ stimulation alone (Figure 2A). In contrast, hippuric acid significantly reduced the production of the anti-inflammatory cytokine IL-10 in the LPS+IFNγ condition (Figure 2A). To further evaluate the activation state of macrophages, we analyzed the surface expression of macrophage activation markers using flow cytometry. Hippuric acid pre-treatment followed by LPS+IFNγ stimulation significantly increased the expression of MHC-I and CD40 compared to LPS+IFNγ alone (Figure 2B). These results indicate that hippuric acid enhances macrophage activation, amplifying both the production of pro-inflammatory cytokines and the expression of co-stimulatory molecules, which are critical for sustaining inflammation.

**Figure 2:**
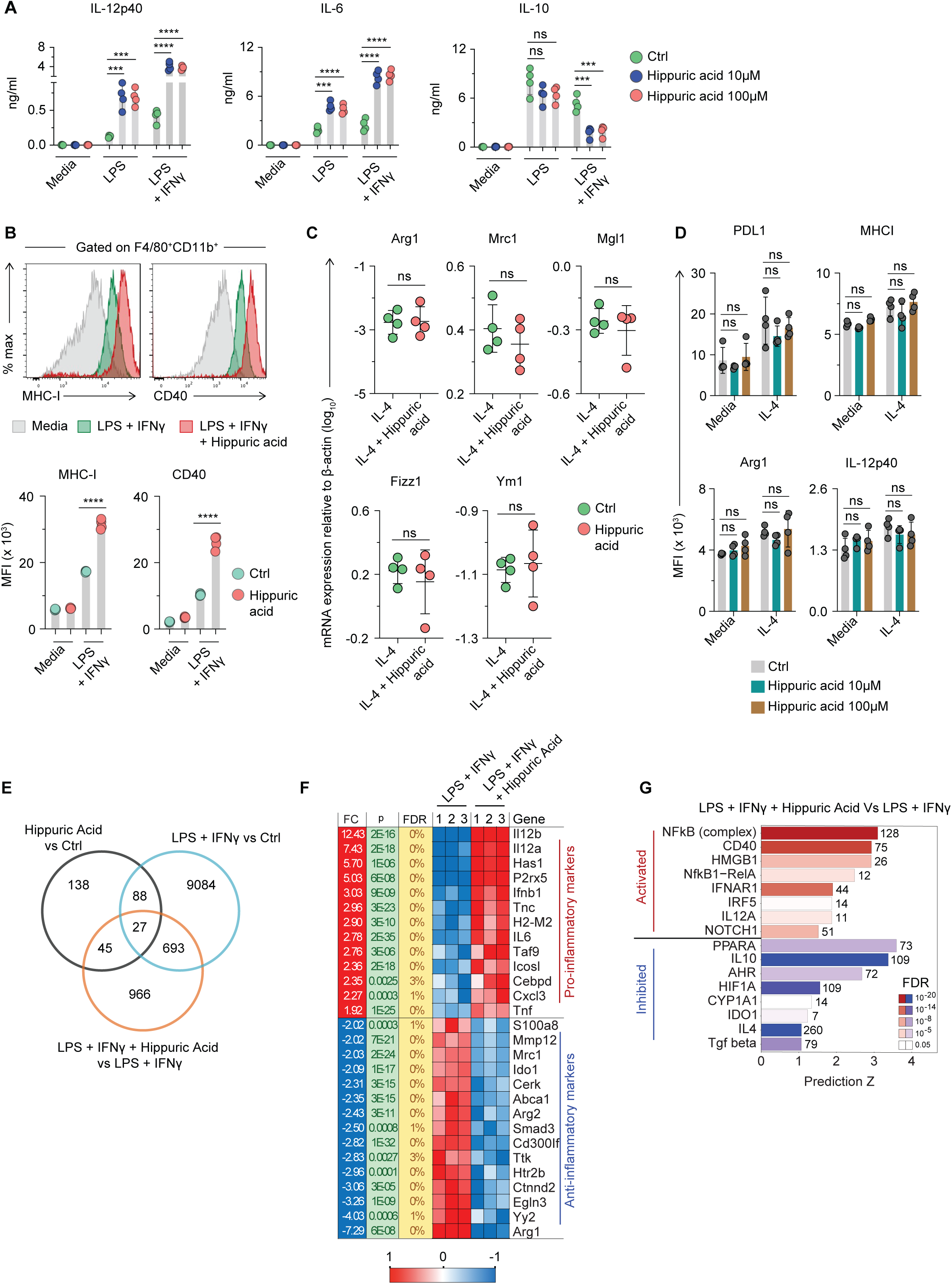
Hippuric acid promotes M1-like macrophage polarization but showed no effect on M2-like polarization. (A) ELISA showing measurements of IL-12p40, IL-6, and IL-10 production by bone marrow derived macrophages (BMDMs) pre-treated with hippuric acid (10µM or 100µM) for 1hr and then stimulated with LPS (100ng/ml) or LPS plus IFNγ (100ng/ml) for 8hr. n=4. (B) Flow cytometry analyses on BMDMs treated as in (A) for surface expression of activation markers MHC-I and CD40 shown as mean fluorescent intensity (MFI). Histogram shows MFI of MHC-I and CD40 on BMDM. n=4. (C) RT-PCR analysis for relative expression of indicated immunosuppressive genes in BMDMs pre-treated with hippuric acid (10µM) for 1hr and then stimulated with IL-4 (10ng/ml) for 8hr. n=4. mRNA relative expression is shown compared to housekeeping gene β-actin. (D) Flow cytometry analyses for surface expression of indicated markers on BMDMs pre-treated with hippuric acid (10µM or 100µM) for 1hr and then stimulated with IL-4 (10ng/ml) for 8hr. Data are presented as mean fluorescent intensity (MFI). n=4. (E-G) RNAseq analysis of BMDMs treated as in (A) for (E) Venn diagram showing overlap of differentially expressed genes comparing hippuric acid vs control (left top circle), M1-like (LPS+IFNγ) vs control (right top circle), and M1-like + hippuric acid vs M1-like (bottom circle), (F) heatmap showing differential gene expression of indicated RNA transcripts for immune-related genes of interest. (G) Ingenuity Pathway Analysis (IPA) of activated (red bars) and inhibited (blue bars) regulators. Differentially expressed genes with FDR <5% were used. n=3. In (A), (B), and (D) *P* values were determined by one-way ANOVA with post hoc multiple comparisons. In (C), *P* values were determined by two-tailed Student’s *t* tests. Data are presented as means ± SD.

Given the pro-inflammatory effects of hippuric acid on M1-like polarization, we next examined whether hippuric acid influences anti-inflammatory responses in M2-like macrophages. To test this, BMDMs were pre-treated with hippuric acid and stimulated with IL-4, a canonical inducer of M2-like polarization, for 8 hours. RT-PCR analysis revealed no significant changes in the expression of M2-like-associated anti-inflammatory genes, including Arg1, Mrc1, Mgl1, Fizz1, and Ym1, when comparing IL-4 stimulation with or without hippuric acid (Figure 2C). Similarly, flow cytometry analysis revealed no significant changes in the surface expression of either anti-inflammatory markers (PDL1 and Arg1) or pro-inflammatory markers (MHCI and IL-12p40) under these conditions (Figure 2D). These findings suggest that hippuric acid selectively enhances M1-like pro-inflammatory responses without significantly impacting M2-like macrophage polarization.

To gain deeper insights into the molecular mechanisms underlying hippuric acid-mediated pro-inflammatory responses, we performed RNA-seq analysis on BMDMs pre-treated with hippuric acid and stimulated with LPS+IFNγ. Hippuric acid treatment alone resulted in significant changes in the expression of 298 genes (FDR <5%), many of which were associated with inflammatory responses (Figure 2E). Ingenuity Pathway Analysis (IPA) revealed activation of pathways related to pro-inflammatory signaling, including IL-6 signaling, iNOS signaling, and production of nitric oxide and reactive oxygen species in macrophages, while pathways associated with immunosuppression, such as phagosome formation, PPAR signaling, and the antioxidant action of vitamin C, were significantly inhibited (Supplementary Figure 2A). When combined with M1-like stimulation (LPS+IFNγ), hippuric acid significantly altered the expression of 1,731 genes (FDR <5%) compared to the M1-like stimulated condition (Figure 2E). Venn diagram analysis revealed a substantial overlap in differentially expressed genes between M1-like stimulated macrophages and those exposed to hippuric acid, underscoring its amplifying effect on M1-like polarization (Figure 2E). A heatmap of selected genes demonstrated that hippuric acid increased the expression of pro-inflammatory genes, including Il12b, Il6, H2-M2, and Tnf, while suppressing anti-inflammatory genes such as Arg1, Smad3, Ido1, and Mrc1 (Figure 2F). IPA of upstream regulators further revealed activation of pro-inflammatory regulators, including NF-κB (complex), CD40, IFNAR1, and IL-12A, alongside inhibition of anti-inflammatory regulators, such as IL-10, AHR, IL-4, and TGF-β (Figure 2G). These transcriptomic findings align with the heightened pro-inflammatory cytokine production and activation marker expression. Together, these results suggest that hippuric acid enhances M1-like polarization by activating pro-inflammatory transcriptional programs while concurrently suppressing pathways involved in immunosuppression and tolerance.

### Hippuric acid increased pro-inflammatory responses to MyD88-dependent TLR ligands but not TRIF-dependent TLR3 ligand

Our initial observation that hippuric acid enhanced pro-inflammatory responses to LPS (a TLR4 ligand) but not IL-4 suggested a specific role for hippuric acid in potentiating TLR signaling. To investigate this further, BMDMs were stimulated with a panel of TLR ligands, including Pam3CSK4 (TLR1/2), LTA and Zymosan (TLR2), Poly(I:C) (TLR3), R848 (TLR7/8), and CpG (TLR9), with or without IFNγ. BMDMs were first pre-treated with hippuric acid (10 µM or 100 µM) and then stimulated with TLR ligands. ELISA analysis of culture supernatants revealed that hippuric acid enhanced the production of IL-12p40 and IL-6 while decreasing IL-10 in response to MyD88-dependent TLR ligands (Pam3CSK4, LTA, Zymosan, R848, and CpG) (Figure 3A-F). Interestingly, hippuric acid had no effect on cytokine production in response to the TRIF-dependent TLR3 ligand Poly(I:C) stimulation (Figure 3D). These results demonstrate that hippuric acid selectively amplifies MyD88-dependent TLR pro-inflammatory responses, while leaving TRIF-dependent activation unaffected, highlighting its specificity as a modulator of inflammatory responses.

**Figure 3:**
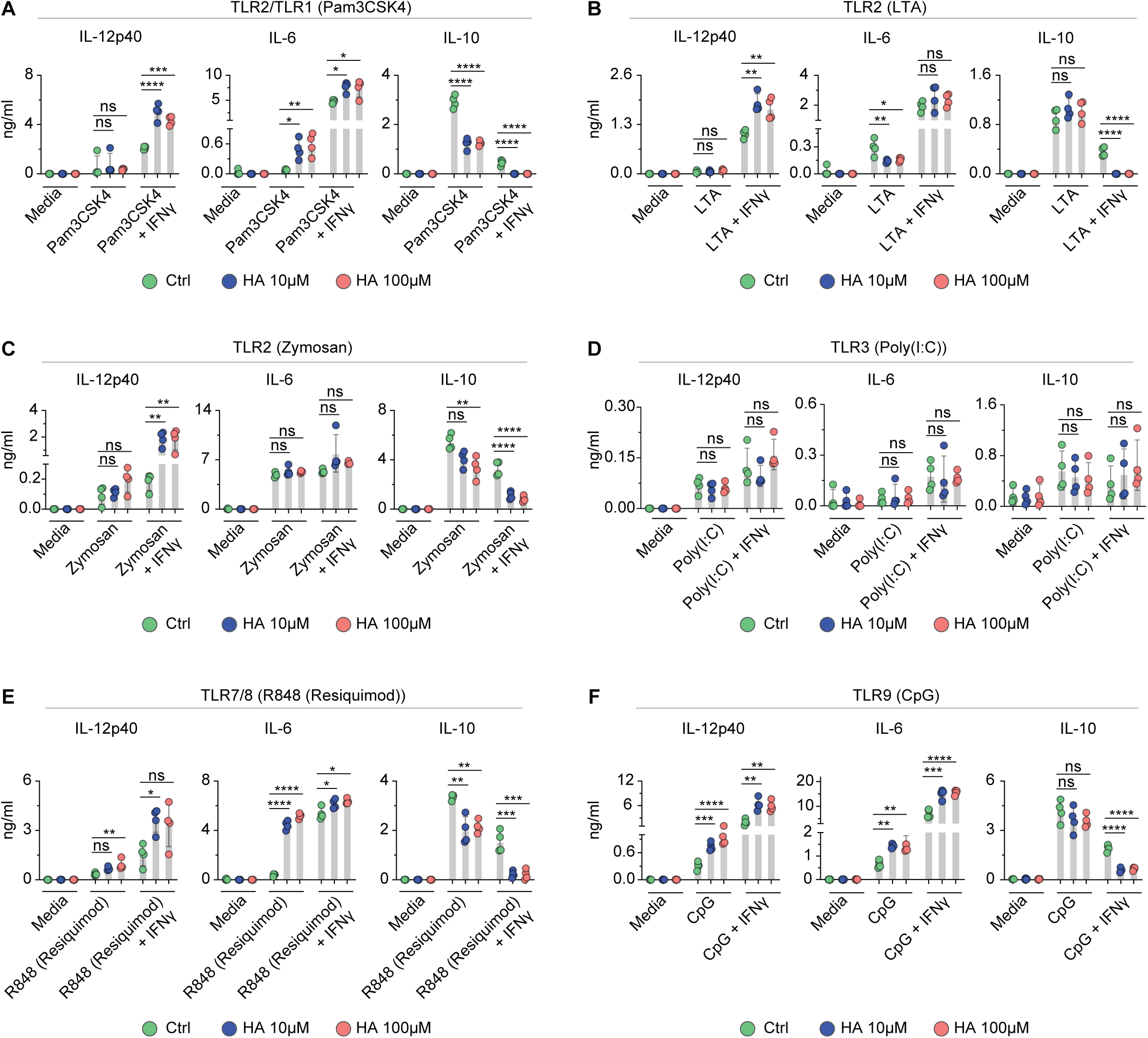
Hippuric acid increases pro-Inflammatory responses to MyD88-dependent TLR ligands but not TRIF-dependent TLR3 ligand. (A-F) ELISA showing measurements of IL-12p40, IL-6, and IL-10 production by bone marrow derived macrophages (BMDMs) pre-treated with hippuric acid (10µM or 100µM) for 1hr and then stimulated with the below Toll-like receptor (TLR) ligands for 8hr. n=4. (A) TLR2/TLR1 (Pam3CSK4) (B) TLR2 (LTA) (C) TLR2 (Zymosan) (D) TLR3 (Poly(I:C)) (E) TLR7/8 (R848 (Resiquimod)) (F) TLR9 (CpG) In (A-F) *P* values were determined by one-way ANOVA with post hoc multiple comparisons. Data are presented as means ± SD.

### MyD88 was required for hippuric acid driven pro-inflammatory responses *in vitro* and *in vivo*

To determine whether the pro-inflammatory effects of hippuric acid are dependent on MyD88, we conducted RNA-seq analysis of BMDMs pre-treated with hippuric acid and stimulated with M1(LPS+IFNγ). IPA identified activation of key regulators in the MyD88 pathway, including TICAM1, TRAF6, and IRAK4, as well as TLR-specific regulators such as TLR1/2, TLR4, TLR7/8, and TLR9 (Figure 4A). These data prompted us to test the requirement of MyD88 for hippuric acid-driven pro-inflammatory responses. We generated BMDMs from wild-type and MyD88 knockout (KO) mice to directly test the requirement of MyD88 in hippuric acid-driven responses. BMDMs were pre-treated with hippuric acid followed by stimulation with M1-like (LPS+IFNγ). ELISA analysis of culture supernatants showed that hippuric acid significantly enhanced IL-12p40 and IL-6 production in WT BMDMs stimulated with M1(LPS+IFNγ) but failed to do so in MyD88-deficient BMDMs (Figure 4B, Supplementary Figure 2B). Similarly, flow cytometry analysis revealed that hippuric acid strongly upregulated macrophage activation markers, including CD86, MHC-I, and CD40, in wild-type BMDMs, but had minimal effects on MyD88-deficient macrophages, particularly for CD86 expression. (Figure 4C, Supplementary Figure 2C). These findings establish MyD88 as a critical mediator of hippuric acid-induced cytokine production and macrophage activation.

**Figure 4:**
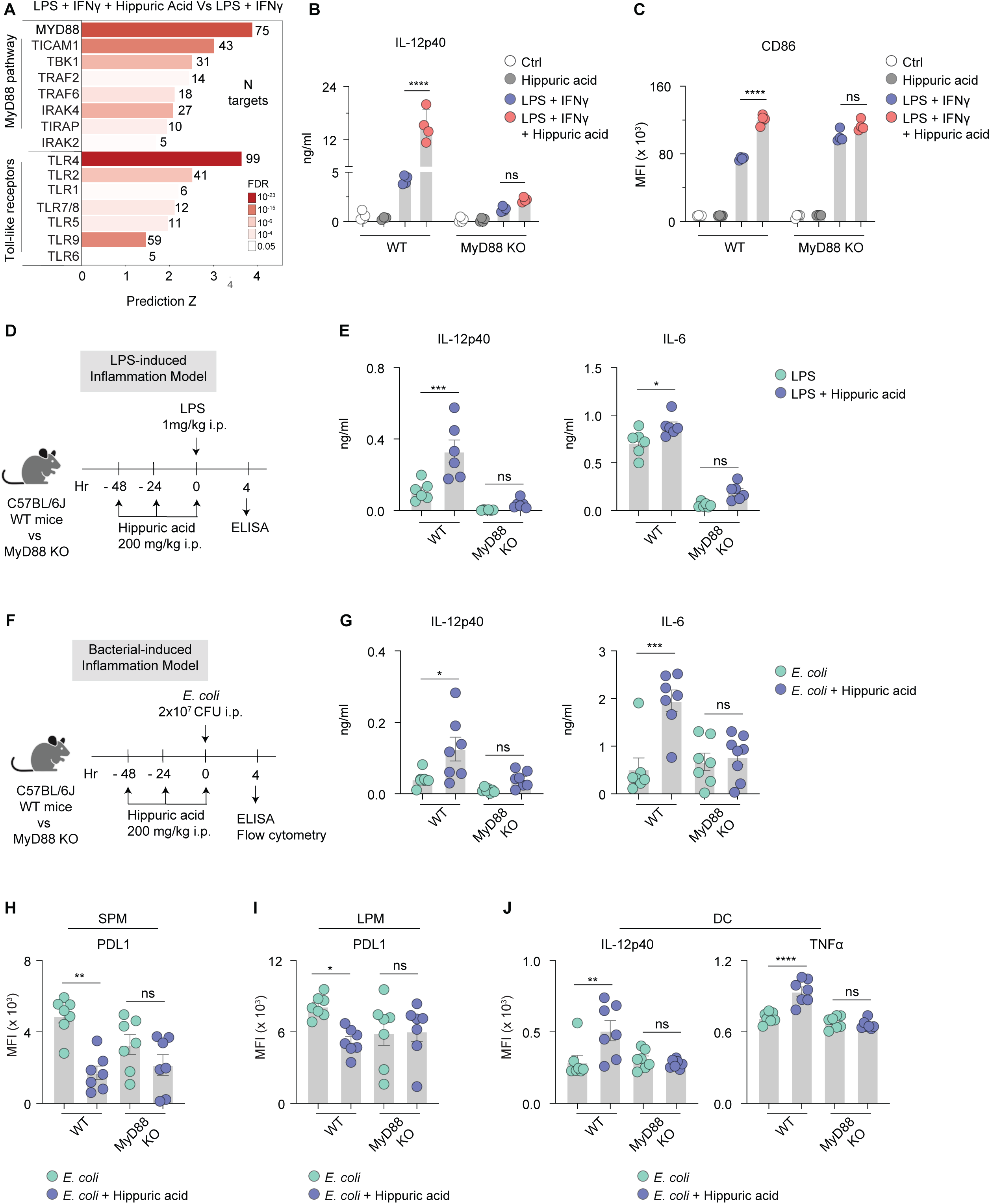
MyD88 is required for hippuric acid driven pro-inflammatory responses *in vitro* and *in vivo*. (A) Ingenuity Pathway Analysis (IPA) of RNAseq for activated (red bars) regulators on bone marrow derived macrophages (BMDMs) pre-treated with hippuric acid (10µM) for 1hr and then stimulated with LPS (100ng/ml) plus IFNγ (100ng/ml) for 8hr. Regulators of interest are grouped under either the MyD88 pathway or the Toll-like receptor family. Differentially expressed genes passing FDR <5% were used. n=3. (B) ELISA showing measurements of IL-12p40 production by BMDMs obtained from wild type or MyD88 KO mice and pre-treated with hippuric acid (10µM) for 1hr and then stimulated with LPS (100ng/ml) plus IFNγ (100ng/ml) for 8hr. n=4. (C) Flow cytometry analyses on BMDMs treated as in (B) showing expression levels of CD86 as mean fluorescent intensity (MFI). n=4. (D) Schematic of LPS-induced inflammation model showing time intervals for administering hippuric acid intraperitoneally and readouts of the study in wild type or MyD88 KO mice. (E) ELISA on peritoneal lavage obtained from mice in (D) for measurements of pro-inflammatory cytokines IL-12p40 and IL-6. n=6 mice per group. (F) Schematic of *E. coli* infection model showing time intervals for administering hippuric acid intraperitoneally and readouts of the study in wild type or MyD88 KO mice. (G) ELISA on peritoneal lavage obtained from mice in (F) for measurements of pro-inflammatory cytokines IL-12p40 and IL-6. n=7 mice per group. (H-J) Flow cytometry analyses of CD45+ immune cells in the peritoneum obtained from mice in (F) showing expression levels PDL1 on (H) small peritoneal macrophages (SPM) and (I) large peritoneal macrophages (LPM), and pro-inflammatory cytokines IL-12p40 and TNFα on (J) dendritic cells (DCs). n=7 mice per group. In (B-C), *P* values were determined by two-way ANOVA with post hoc multiple comparisons. In (E), (G), and (H-J), *P* values were determined by one-way ANOVA with post hoc multiple comparisons. Data are presented as means ± SD.

To validate these findings *in vivo*, we employed LPS-induced inflammation and *E. coli* infection models in wild-type and MyD88-deficient mice. In the LPS model, wild-type mice pre-treated with hippuric acid exhibited significantly increased levels of IL-12p40 and IL-6 in the peritoneal lavage compared to vehicle-treated controls, while MyD88-deficient mice showed no such increase (Figure 4D-E). Similarly, in the *E. coli* infection model, hippuric acid enhanced IL-12p40 and IL-6 production in wild-type mice but not in MyD88-deficient mice (Figure 4F-G). Flow cytometry analysis on the peritoneal lavage fluid revealed that hippuric acid decreased PDL1 expression on small (SPMs) and large peritoneal macrophages (LPMs) and enhanced IL-12p40 and TNFα production by dendritic cells (DCs) in wild-type mice, with no changes observed in MyD88-deficient mice (Figure 4H-J). Collectively, these results demonstrate that MyD88 is required for hippuric acid to amplify pro-inflammatory cytokine production and innate immune activation during inflammation and infection.

### Hippuric acid drove cholesterol biosynthesis and lipid remodeling in macrophages

Given the established interplay between TLR-MyD88 signaling, lipid metabolism, and pro-inflammatory responses in macrophages ^22–24^, we next investigated whether hippuric acid modulates macrophage lipid remodeling. IPA of RNA-seq on BMDMs stimulated with M1-like (LPS+IFNγ) in the presence of hippuric acid revealed the activation of pathways related to lipid remodeling and heightened pro-inflammatory responses, including the superpathway of cholesterol biosynthesis, cholesterol biosynthesis I, mevalonate pathway I, and HMGB1 signaling, while anti-inflammatory pathways such as phagosome formation, PPAR signaling, and STAT3 signaling were significantly inhibited (Figure 5A). To further characterize lipid metabolism, we queried the expression of lipid-related genes using the LIPID MAPS proteome database (LMPD), which contains a curated list of lipid-associated genes and proteins ^27^. Out of 1504 entries in the LMPD, 1067 were detected in RNA-seq data. A heatmap of selected genes passing FDR <5% and fold change >1.5 demonstrated that hippuric acid increased the expression of genes involved in cholesterol biosynthesis, including Hmgcs1, Sqle, Srebf2, and Dhcr24, while downregulating genes associated with cholesterol export, such as Abca1 and Soat2 (Figure 5B).

**Figure 5:**
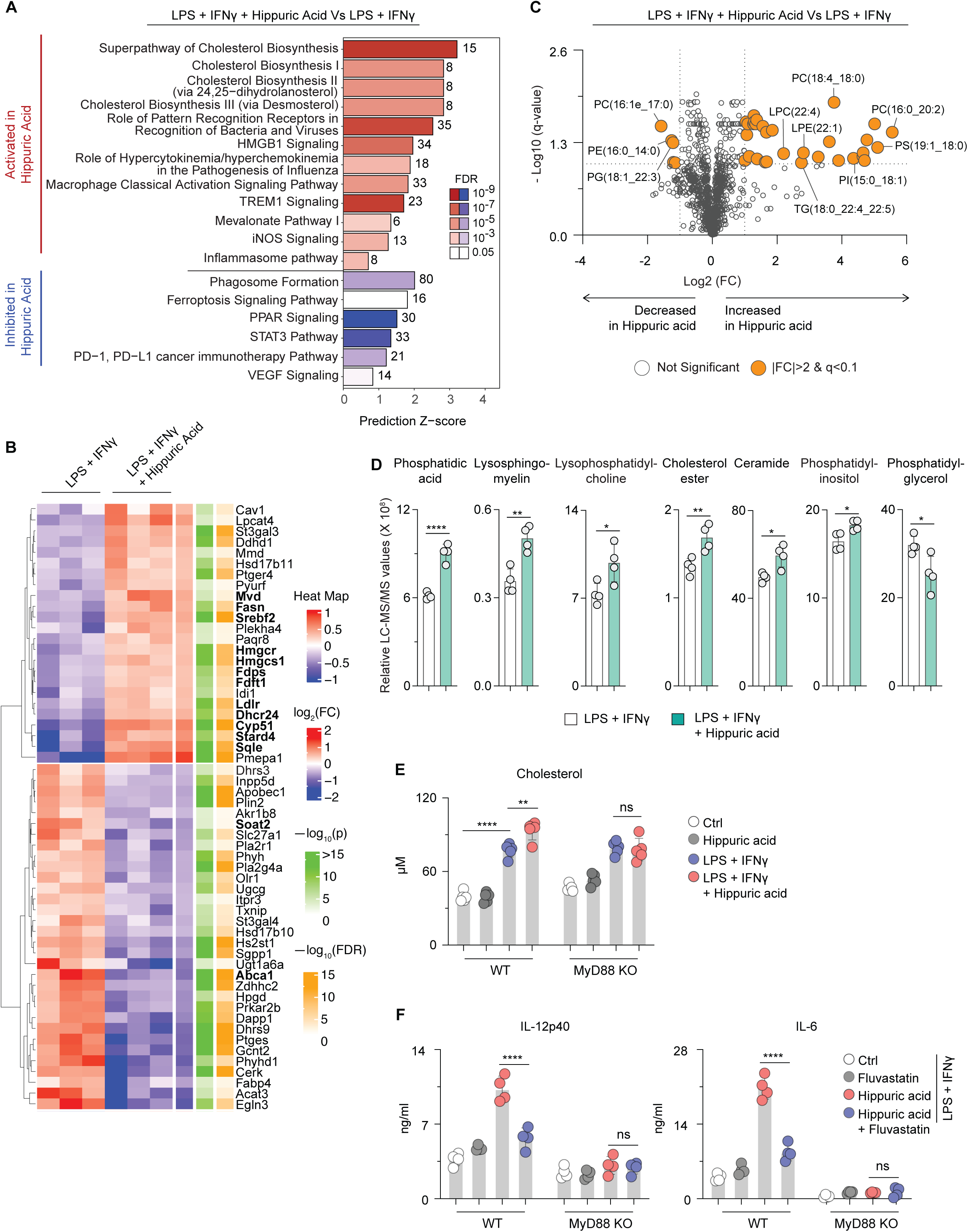
Hippuric acid drives cholesterol biosynthesis and lipid remodeling in macrophages. (A) Ingenuity Pathway Analysis (IPA) of RNAseq for activated (red bars) and inhibited (blue bars) pathways on bone marrow derived macrophages (BMDMs) pre-treated with hippuric acid (10µM) for 1hr and then stimulated with LPS (100ng/ml) plus IFNγ (100ng/ml) for 8hr. Differentially expressed genes passing FDR <5% were used. n=3. (B) RNAseq analysis of BMDMs treated as in (A) for heatmap showing differential gene expression of indicated RNA transcripts of lipid metabolism using the LIPID MAPS proteome database (LMPD). Differentially expressed genes passing FDR <5% and fold change >1.5 were used. n=3. (C) Lipidomics analysis of BMDMs treated as in (A) for volcano plot showing changes in individual lipid species. Lipidomics was performed using LC-MS/MS. n=4. (D) Lipidomics analysis of BMDMs treated as in (A) for changes in lipid classes shown as bar graphs. n=4. (E) Colorimetric assay for cholesterol quantitation in BMDMs obtained from wild type or MyD88 KO mice and pre-treated with hippuric acid (10µM) for 1hr and then stimulated with LPS (100ng/ml) plus IFNγ (100ng/ml) for 8hr. n=4. (F) ELISA showing measurements of IL-12p40 and IL-6 production by BMDMs obtained from wild type or MyD88 KO mice and pre-treated with hippuric acid (10µM) or fluvastatin (10µM) for 1hr and then stimulated with LPS (100ng/ml) plus IFNγ (100ng/ml) for 8hr. n=4. In (C), significant change was defined as |FC| > 2; *q* < 0.1 (Benjamini-Hochberg FDR-adjusted *P* value). In (D), *P* values were determined by two-tailed Student’s *t-tests*. In (E-F), *P* values were determined by two-way ANOVA with post hoc multiple comparisons. Data are presented as means ± SD.

Prompted by these findings, we assessed whether hippuric acid broadly perturbs lipid metabolism by performing lipidomics analysis on BMDMs pre-treated with hippuric acid and stimulated with M1-like (LPS+IFNγ). Lipids were extracted 24 hours after stimulation and analyzed via liquid chromatography-tandem mass spectrometry (LC-MS/MS). Consistent with prior studies, M1-like macrophages displayed significant alterations in lipid composition compared to controls, including the accumulation of distinct triglyceride (TG) species and reductions in cholesterol ester (ChE) and lysophosphatidylcholine (LPC) species (Supplementary Figure 3A-B). Addition of hippuric acid drove changes in lipid composition in M1-polarized macrophages, with significant increases in phospholipids and neutral lipids. For example, lipid species such as PC(16:0_20:2), PS(19:1_18:0), and PI(18:0_17:0) increased >27-fold, and LPE(22:1), TG(18:0_22:4_22:5), and LPC(22:4) increased >4-fold. Only a few lipid species, such as PC(16:1e_17:0) and PE(16:0_14:0), were downregulated with <3-fold changes (Figure 5C). Analysis of lipid classes revealed significant increases in phosphatidic acid, lysosphingomyelin, lysophosphatidylcholine, cholesterol ester, ceramide, and phosphatidylinositol, while phosphatidylglycerol levels decreased (Figure 5D).

Cholesterol accumulation is a hallmark of the M1-like macrophage phenotype, facilitating TLR-MyD88 signaling and driving pro-inflammatory cytokine production ^17,26^. To determine whether hippuric acid-mediated cholesterol synthesis depended on MyD88 signaling, we performed a colorimetric assay to measure total cholesterol levels in BMDMs pre-treated with hippuric acid and stimulated with LPS+IFNγ. Hippuric acid significantly increased cholesterol levels in WT BMDMs stimulated with M1-like (LPS+IFNγ) compared with unstimulated controls. However, this increase was absent in MyD88-deficient BMDMs, indicating that cholesterol accumulation in response to hippuric acid requires MyD88 signaling (Figure 5E). We next investigated whether cholesterol accumulation is required for the pro-inflammatory effects of hippuric acid. We used fluvastatin, an inhibitor of cholesterol biosynthesis, in LPS+IFNγ-treated macrophages. Fluvastatin significantly reduced the production of IL-12p40 and IL-6 in WT BMDMs treated with hippuric acid, but this was unchanged in MyD88-deficient BMDMs (Figure 5F). These findings demonstrate that cholesterol accumulation is a critical mediator of the pro-inflammatory effects of hippuric acid, linking lipid metabolism to TLR-MyD88-driven macrophage activation. Taken together, these results establish that hippuric acid amplifies TLR-MyD88 signaling through enhanced cholesterol biosynthesis and lipid remodeling in macrophages.

### A pro-inflammatory effect of hippuric acid was observed in human macrophages

To extend our findings to human settings, we tested whether hippuric acid potentiates pro-inflammatory responses in human macrophages. Human monocyte-derived macrophages (HMDMs) were pre-treated with hippuric acid for 1 hour and stimulated with LPS+IFNγ for 8 hours. RT-PCR analysis revealed significant upregulation of pro-inflammatory cytokines, including IL-6, IL-1β, and TNFα, along with increased expression of cholesterol biosynthesis genes such as HMGCS1, SREBF2, and SQLE, while anti-inflammatory cytokine IL-10 was notably decreased (Figure 6A-B). Consistently, ELISA analysis showed that hippuric acid enhanced IL-6 production and decreased IL-10 levels in response to M1-like (LPS+IFNγ) stimulation (Figure 6C). These results demonstrate that hippuric acid amplifies pro-inflammatory responses in human macrophages, functioning similarly to its effects in murine macrophages.

**Figure 6:**
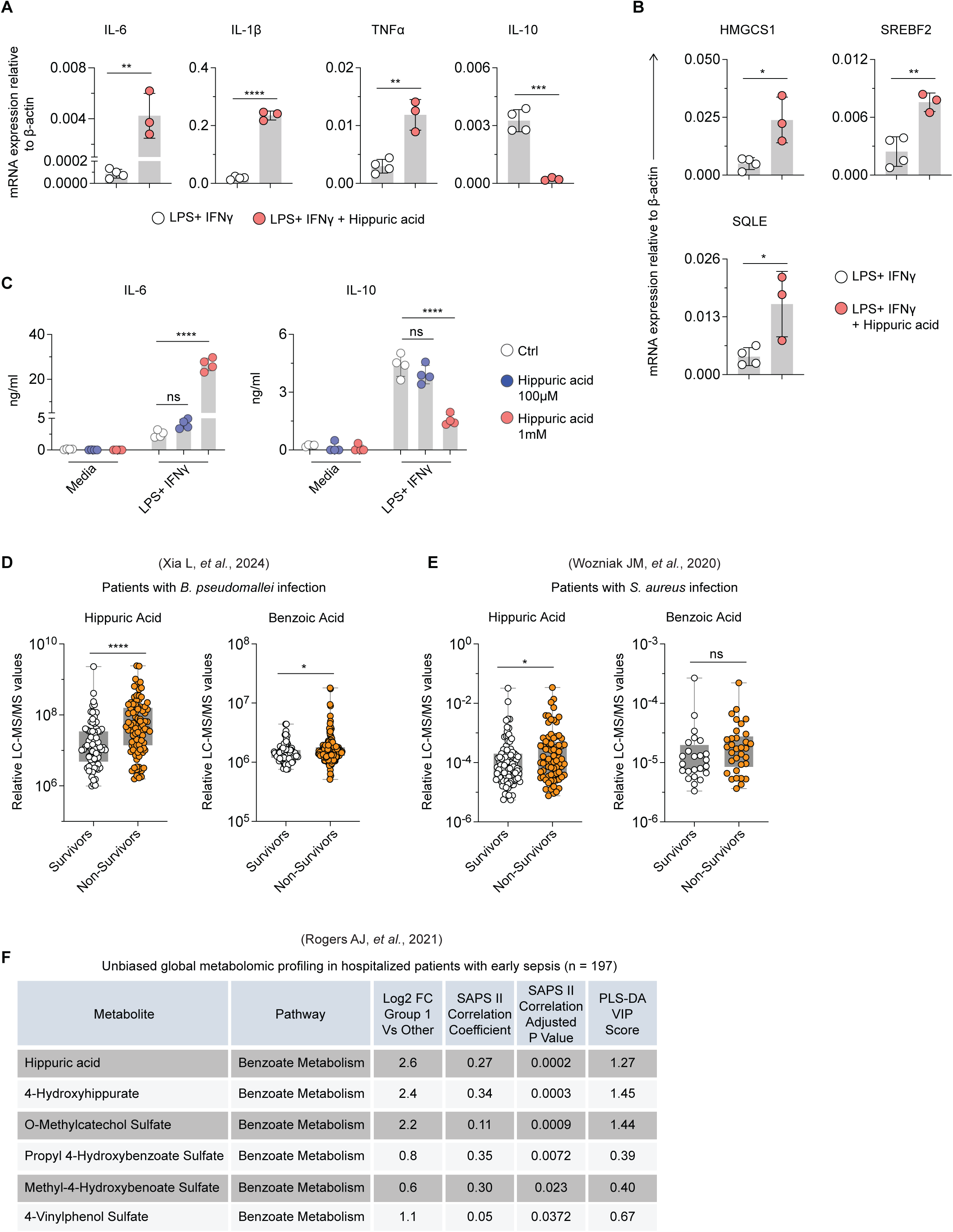
Levels of hippuric acid correlates with disease severity and increased mortality in patients with sepsis. (A-B) RT-PCR analysis for relative expression of immunoregulatory cytokines including IL-6, IL-1β, TNFα, and IL-10, and cholesterol synthesis pathway genes including HMGCS1, SREBF2, and SQLE in human monocyte derived macrophages (HMDMs) pre-treated with hippuric acid (100µM) for 1hr and then stimulated with LPS (100ng/ml) plus IFNγ (100ng/ml) for 8hr. n=3-4. mRNA relative expression is shown compared to housekeeping gene β-actin. *P* values were determined by two-tailed Student’s *t-tests*. Data are presented as means ± SD. (C) ELISA showing measurements of immunoregulatory cytokines IL-6 and IL-10 production by HMDMs treated as in (A). n=4. *P* values were determined by two-way ANOVA with post hoc multiple comparisons. Data are presented as means ± SD. (D-E) Box-and-whisker plot for relative abundance of hippuric acid and benzoic acid in survivors versus non-survivors of patients with (D) *B. pseudomallei* and (E) *S. aureus* infection. *P* values were determined by a Mann-Whitney nonparametric test for comparisons of relative abundance values between survivor versus non-survivor patients. (F) Comparison of plasma levels of hippuric acid and metabolites within benzoic metabolism pathway in patients with early sepsis (n=197) ranked by PLS-DA VIP score. Metabolite comparisons include Group 1 (high mortality group) relative to all other patients and association with severity of illness (SAPS II score). SAPS II P value is for Pearson’s correlation after adjustment for multiple comparisons. Abbreviations: FC, fold change; SAPS, Simplified Acute Physiology Score; PLS-DA VIP, Partial Least Squares Discriminant Analysis Variable Importance in Projection.

### Levels of hippuric acid correlated with disease severity and increased mortality in patients with sepsis

The gut microbiota plays a critical role in sepsis progression by modulating immune responses and systemic inflammation ^28,29^. Sepsis disrupts gut barrier integrity, enabling bacterial translocation into circulation and exacerbating inflammatory responses, while also altering microbiome composition and microbial metabolite levels. Microbial metabolites, in turn, regulate key immune processes, such as macrophage function, which shape susceptibility to sepsis. Despite growing recognition of the role of microbiota, fundamental questions remain about the specific contributions of microbial metabolites to sepsis pathophysiology. Profiling these metabolites could reveal biomarkers for early diagnosis, disease severity, and prognosis, while identifying novel therapeutic targets for restoring immune balance and improving patient outcomes.

Given that our studies implicated hippuric acid in potentiating pro-inflammatory responses during bacterial infection, we investigated its clinical relevance in sepsis patients. A prior study by *Xia L et al* identified distinct metabolic perturbations during infections with *Burkholderia pseudomallei, Escherichia coli, Staphylococcus aureus,* or *Klebsiella pneumoniae*, along with uninfected healthy controls and patients with suspected infections but negative blood cultures ^30^. Examining hippuric acid and its precursor benzoic acid, we found no significant differences in their levels across the groups (Supplementary Figure 4A-B). However, when comparing survivors and non-survivors in *B. pseudomallei* infection, hippuric acid and benzoic acid levels were significantly elevated in non-survivors compared to survivors (Figure 6D). A separate study by *Wozniak JM et al* analyzed serum metabolites in patients with *S. aureus* infection and healthy controls ^31^. Similar to *B. pseudomallei* infection, hippuric acid levels were significantly higher in non-survivors compared to survivors of *S. aureus* infected patients, whereas benzoic acid levels showed no difference (Figure 6E). Collectively, these findings indicate that hippuric acid negatively correlates with survivorship in sepsis, underscoring its potential as a marker of disease severity.

Early sepsis is often associated with heightened inflammation, organ failure, and high mortality ^32^. We hypothesized that elevated levels of hippuric acid might correlate with increased mortality risk in early sepsis patients. Using data from *Rogers AJ et al* ^33^, we analyzed the abundance of hippuric acid and metabolites within the benzoate metabolism pathway in critically ill patients with varying mortality risks. Patients in the high-risk group for 60-day mortality (Group 1) exhibited significantly higher levels of hippuric acid and metabolites within benzoate metabolism pathway (e.g., 4-hydroxyhippurate, o-methylcatechol sulfate, etc.) compared to others in the low- or intermediate-risk groups (Figure 6F). Taken together, these results identify hippuric acid as a marker of systemic inflammation, disease severity, and mortality in sepsis patients, offering potential utility as a prognostic biomarker and therapeutic target.

## Discussion

Our findings establish hippuric acid as a potent microbial-derived modulator of inflammation, linking gut microbial metabolism, innate immune activation, and lipid remodeling. Prior studies have largely characterized hippuric acid as a biomarker of diet, kidney function, and metabolic disorders, with little focus on its direct immunomodulatory effects ^10–13,34^. Our study challenges this passive role by demonstrating that hippuric acid is a bioactive metabolite capable of potentiating pro-inflammatory responses. Mechanistically, hippuric acid promoted macrophage activation via TLR-MyD88 signaling and drove lipid metabolism—particularly cholesterol biosynthesis—to sustain inflammation. These findings expand the functional repertoire of microbial metabolites beyond immunoregulatory short-chain fatty acids (SCFAs), tryptophan metabolites, and bile acid derivatives ^3–8^, providing new insights into how gut microbial metabolism shapes host immunity.

A key finding of our study is the selective enhancement of M1-like macrophage polarization by hippuric acid, without affecting M2-like polarization. While microbial metabolites like butyrate and propionate are well-documented for their anti-inflammatory effects through HDAC inhibition and GPCR activation ^35–37^, emerging evidence suggests that certain metabolites can drive pro-inflammatory responses. For example, our prior work and work by others showed that TMAO, derived from microbial metabolism of choline, is linked to macrophage-driven pro-inflammatory responses in atherosclerosis and cancer ^3,9^. In this study we identified hippuric acid, an aromatic microbial metabolite, acts as a driver of pro-inflammatory responses, likely reinforcing immune responses during infection or inflammatory settings. Moreover, its ability to amplify macrophage responses across multiple TLR ligands suggests that hippuric acid functions as an inflammatory sensitizer, providing co-stimulatory signals that enhance immune activation. This parallels host-derived TCA cycle metabolites, such as succinate, which boosts TLR signaling via SUCNR1 engagement ^38^, and itaconate, which enhances macrophage activation by modulating NF-κB signaling ^39^. Like these metabolites, hippuric acid appears to prime macrophages for heightened inflammatory responses, particularly during infection or acute inflammatory stimulation.

Mechanistically, we demonstrated that hippuric acid potentiated pro-inflammatory responses through MyD88-dependent TLR signaling. This positions hippuric acid within the broader context of host-microbe interactions, where microbial metabolites modulate host immunity by influencing conserved pattern recognition receptor (PRR) pathways ^4,40^. While our findings show that hippuric acid amplifies TLR signaling responses, it remains unclear whether it acts as a direct TLR ligand. However, our observations align with emerging evidence that microbial metabolic cues can fine-tune TLR signaling dynamics. For example, SCFAs, indole derivatives, and bile acid derivatives have been shown to modulate TLR signaling ^4,40–42^, suggesting that microbial metabolites play an active role in shaping innate immune responses.

Interestingly, hippuric acid potentiated inflammatory responses to most TLRs, including TLR1, TLR2, TLR4, TLR7/8, and TLR9, but not to TLR3. This discrepancy likely reflects the distinct signaling mechanisms of TLR3 compared to other TLRs. Most TLRs signal through the MyD88-dependent pathway, which activates NF-κB and MAPK pathways to induce the production of pro-inflammatory cytokines such as IL-6, TNFα, and IL-12p40 ^43,44^. However, TLR3 uniquely signals via the TRIF adaptor protein, driving type I interferon responses, which involve IRF3 phosphorylation and production of IFN-β ^45,46^. Unlike the pro-inflammatory cytokines induced by MyD88 signaling, type I interferons can have pro-inflammatory and anti-inflammatory effects ^47^, which might explain why hippuric acid selectively potentiates inflammatory cytokine production while leaving TRIF/IFN-I pathways unaltered. Furthermore, MyD88-dependent signaling relies heavily on protein-protein interactions mediated by hydrophobic and aromatic domains, providing a potential mechanism for the selective action of hippuric acid ^44,48–50^. As an aromatic microbial metabolite containing a benzoyl moiety, hippuric acid may facilitate MyD88 recruitment by engaging with aromatic and hydrophobic binding pockets within the intracellular domains of TLRs, thereby enhancing signal transduction. Such interactions could underlie the heightened inflammatory cytokine production observed in macrophages exposed to hippuric acid, providing a mechanistic explanation for its selective amplification of MyD88-driven pathways.

One of the most intriguing findings of this study is the role of lipid remodeling and cholesterol biosynthesis in hippuric acid-driven macrophage activation. Lipid metabolism is increasingly recognized as a key determinant of macrophage function, influencing phagocytosis, antigen presentation, and cytokine production ^51,52^. Our transcriptomic and lipidomic analyses revealed that hippuric acid promoted cholesterol biosynthesis and lipid accumulation in macrophages, a feature commonly associated with inflammatory macrophage activation in atherosclerosis and metabolic disorders ^53^. This is consistent with previous studies showing that TLR activation is tightly linked to lipid metabolism, as TLR signaling reshapes macrophage lipid composition to support inflammatory responses ^24,25,54,55^. TLRs, especially TLR4, are known to localize in cholesterol-rich lipid rafts, where receptor clustering and signal transduction occur ^56^. It is possible that hippuric acid alters lipid raft composition or enhances cholesterol-dependent membrane microdomain formation, thereby amplifying TLR signaling and macrophage activation. This hypothesis aligns with previous studies showing that lipid metabolism and innate immune activation are tightly linked, and that metabolic rewiring of macrophages can sustain inflammatory responses.

Beyond TLR-MyD88 signaling and lipid remodeling, hippuric acid may influence macrophage responses through interactions with nuclear receptors and metabolic enzymes. Aromatic microbial metabolites are known to modulate nuclear receptors such as the aryl hydrocarbon receptor (AhR), peroxisome proliferator-activated receptors (PPARs), and liver X receptors (LXRs), which link metabolic and immune pathways and macrophage activation ^2,57,58^. As an aromatic compound, hippuric acid may interact with these lipid-sensing transcription factors, affecting inflammatory gene expression and macrophage metabolism. Given the structural similarity to aromatic AhR agonists, hippuric acid could modulate AhR activity, potentially inhibiting its anti-inflammatory effects or competing with classical AhR ligands, thus influencing macrophage polarization. Similar mechanisms have been observed for oxidized lipids and bacterial-derived phenolic metabolites, which disrupt PPARγ activation and promote metabolic inflammation ^59^. As hippuric acid drove cholesterol biosynthesis and lipid accumulation, it may also interfere with LXR signaling, disturbing lipid homeostasis and sustaining inflammation^57^. Future studies are needed to investigate whether hippuric acid directly binds to these receptors or modulates their downstream effects on macrophage function.

Clinically, elevated hippuric acid levels correlated with higher mortality in sepsis patients, underscoring its potential role in inflammatory diseases. Sepsis is characterized by dysregulated immune responses, in which macrophage hyperactivation and excessive cytokine production contribute to systemic inflammation and organ failure ^28,29^. Microbial metabolites such as SCFAs, indoles, and TMAO are now recognized as key immune modulators, influencing macrophage polarization and inflammatory gene expression. Our findings suggest that hippuric acid may serve as a biomarker or therapeutic target for inflammatory conditions where macrophage overactivation drives pathology. Notably, viral infections such as influenza and COVID-19 exhibit similar immune dysregulation with metabolites like tryptophan-derived kynurenines and SCFAs influencing disease severity through macrophage modulation ^60^. Future research should investigate whether targeting hippuric acid production or its downstream signaling pathways could mitigate excessive inflammation in sepsis and other immune-driven disorders.

In conclusion, we have identified hippuric acid as a previously unrecognized microbial metabolite that potentiates macrophage-driven inflammation through TLR-MyD88 signaling and lipid remodeling. More broadly, this work highlights the role of aromatic microbial metabolites in inflammation, suggesting that beyond SCFAs, tryptophan metabolites, and bile acids, gut microbes produce a diverse spectrum of immunomodulatory metabolites that influence host immunity and may provide novel therapeutic targets for inflammatory and infectious diseases.

---

**Supplementary Figure 1: Hippuric acid influences local and systemic inflammation in murine models of *E. coli* infection and LPS-induced inflammation.**

(A-H) Flow cytometry analyses of CD45+ immune cells for percent populations or expression levels of indicated markers. C57BL/6 mice were administered with hippuric acid intraperitoneally and then infected with E. coli for 4hr. CD45+ immune cells were obtained from (A-D) peritoneal lavage, (E) bone marrow, or (F-H) spleen. n=7 mice per group.

(I-K) Flow cytometry analyses of CD45+ immune cells obtained from peritoneal lavage for expression levels of indicated markers. C57BL/6 mice were administered with hippuric acid intraperitoneally and then treated with LPS for 4hr. n=5 mice per group.

*P* values were determined by two-tailed Student’s *t-tests*. Data are presented as means ± SD

**Supplementary Figure 2: Hippuric acid directly enhances pro-inflammatory effects in macrophages.**

(A) RNAseq analysis of bone marrow derived macrophages (BMDMs) treated with hippuric acid for 8hr. Ingenuity Pathway Analysis (IPA) of activated (red circles) and inhibited (blue circles) pathways in hippuric acid treated compared to control macrophages. Differentially expressed genes passing FDR <5% were used. n=3.

(B) ELISA showing measurements of IL-6 production by BMDMs obtained from wild type or MyD88 KO mice and pre-treated with hippuric acid (10µM) for 1hr and then stimulated with LPS (100ng/ml) plus IFNγ (100ng/ml) for 8hr. n=4.

(C) Flow cytometry analyses on BMDMs treated as in (B) showing expression levels of MHCI and CD40 as mean fluorescent intensity (MFI). n=4.

In (B-C), P values were determined by two-way ANOVA with post hoc multiple comparisons. Data are presented as means ± SD.

**Supplementary Figure 3: M1-like polarization accompanies distinct changes in lipid profiles in macrophages.**

(A) Lipidomics analysis of on bone marrow derived macrophages (BMDMs) pre-treated with hippuric acid (10µM) for 1hr and then stimulated with LPS (100ng/ml) plus IFNγ (100ng/ml) stimuli for 8hr. Volcano plot shows changes in individual lipid species. Lipidomics was performed using non-targeted liquid chromatography-tandem mass spectrometry (LC-MS/MS). n=4.

(B) Lipidomics analysis of BMDMs treated as in (A) for changes in lipid classes shown as bar graphs. n=4.

In (A), significant change was defined as |FC| > 2; *q* < 0.1 (Benjamini-Hochberg FDR-adjusted *P* value). In (B), *P* values were determined by two-tailed Student’s *t-tests*. Data are presented as means ± SD.

**Supplementary Figure 4: Levels of hippuric acid are comparable in patients with bacterial infections.**

(A-B) Box-and-whisker plot for relative abundance of (A) hippuric acid and (B) benzoic acid in patients with bacterial infections. *B.pseudomallei (Burkholderia pseudomallei); E. coli (Escherichia coli); K. pneumoniae (Klebsiella pneumoniae); S. aureus (Staphylococcus aureus)*. *P* values were determined by a Kruskal-Wallis test for comparisons of relative abundance values between patients with bacterial infections. Data are presented as means ± SD.

## Materials and Methods

### Animals

Mouse experiments were performed following National Institutes of Health (NIH) guidelines and were approved by the Institutional Animal Care and Use Committee (IACUC) of The Wistar Institute. C57BL6/J (B6), MyD88^-/-^ mice were obtained from Jackson Laboratory and/or were bred at the animal facility of The Wistar Institute. All mice were maintained in a specific pathogen-free barrier facility at The Wistar Institute in accordance with the Association for Assessment and Accreditation of Laboratory Animal Care (AAALAC). Female or male mice at 8-10 weeks old were used in all experiments.

### Generation of primary mouse and human macrophages

Mouse bone marrow derived macrophages (BMDMs) were generated by culturing bone marrow cells with macrophage colony stimulating factor (MCSF) (BioLegend, 576406) as previously described ^7,61^. Briefly, hind legs of 6–8-week-old B6 mice were dissected to harvest femur and tibia and bone marrow cells were isolated by flushing the bone marrow cavity with the help of 27G needle and cold sterile media. Bone marrow cells were cultured in DMEM with10% FBS (Corning, MA, 35-016-CV) and 100 U penicillin and streptomycin antibiotics in the presence of 40ng/ml MCSF (BioLegend, 576406). After 5-7d of differentiation, BMDMs were harvested using ice-cold detachment buffer containing 1XPBS, 2% FBS, and 2mM EDTA (Invitrogen, NY, 15575-038) and used in experiments.

Human monocyte derived macrophages (HMDM) were generated by culturing human monocytes with human MCSF (BioLegend, 574804). Monocytes isolated from PBMCs from healthy donors were obtained from the University of Pennsylvania Human Immunology Core and cultured in DMEM with10% FBS (Corning, MA, 35-016-CV) and 100 U penicillin and streptomycin antibiotics in the presence of 40ng/ml human MCSF. After 5-7d of differentiation, HMDM were harvested using ice-cold detachment buffer and used in experiments.

Additionally, macrophages were polarized to M1-like phenotype by LPS (100ng/ml; Sigma, L2880) or LPS (100ng/ml) plus IFNγ (100ng/ml; BioLegend, 575306), or to M2-like phenotype by IL-4 (10ng/ml; BioLegend, 574304) stimulation for 8hr. Macrophages were stimulated with TLR ligands-Lipoteichoic acid (tlrl-lta; InvivoGen), Pam3CSK4 (tlrl-pms; InvivoGen), Poly (I:C) (tlrl-picw; InvivoGen), Zymosan (tlrl-zyn; InvivoGen), R848 (tlrl-r848; InvivoGen) plus IFNγ and/or Hippuric Acid (HA) (10, 100 uM; Sigma, 112003) for 8 hr. BMDMs were treated with Fluvastatin (100uM; Cayman chemical, 10010334) and Simvastatin (10uM; Cayman chemical, 10010344) for 8 hours following HA (100uM) treatment for 1 hour.

### Enzyme-linked immunosorbent assay (ELISA)

Supernatants from BMDM culture polarized with TLR stimuli were assessed for cytokine production by Enzyme-linked immunosorbent assays (ELISAs). Commercially available kits from Invitrogen were used to measure IL-6 (Invitrogen, 88-7064), IL-12p40 (Invitrogen, 88-7120), and IL-10 (Invitrogen, 88-7105) according to the instructions provided by the manufacturer.

### Reverse transcription polymerase chain reaction (RT-PCR)

Mouse or human macrophage lysates were processed to obtain purified RNA using RNeasy Plus Mini Kit (Qiagen, 74136). iScript™ Reverse Transcription Supermix kit (Bio-Rad, 1708841) was used to reverse transcribe 100ng of RNA to cDNA. RT-PCR reactions were carried out on Applied Biosystems Quant Studio 7 Flex Real-Time PCR system using 1X SYBR Green PCR Master Mix (Applied Biosystems). Primer sequences are listed in Supplementary Table 1. The relative mRNA expression is analyzed compared to the expression of β-actin. Data were analyzed using manufacturer’s instructions.

### *In vivo* treatments

HA (Sigma, 112003) at 200mg/kg was administered via intraperitoneal (i.p.) injection 3x before the acute LPS and E.coli infections. For acute LPS infection, mice were injected with 1mg/kg LPS (Sigma, L2880) via i.p. route and then mice were euthanized after 4 hours.

### *Escherichia coli* peritoneal infection

*Escherichia coli* peritoneal infection was performed as previously described ^62^. Briefly, *E. coli* (ATCC 25922) was cultured in LB medium with 100 µg/ml ampicillin overnight at 37 °C with gentle shaking to an optical density at wavelength 600 nm of 0.5−0.6 to achieve a logarithmic growth phase. C57BL/6J or MyD88 KO mice of 8–12 weeks of age received an intraperitoneal (i.p.) injection of 100 μl PBS with 2 × 10^7^ *E. coli*. Samples were then collected at indicated time points after the infection.

### Flow cytometry

Fluorescent conjugated antibodies used for flow cytometry are listed in the Supplementary Table 2. Spleens were minced with scissors and using the plunger/piston end of the syringe and tissue suspensions were passed through 100μm cell strainers. Bone marrow cells were flushed with complete DMEM. Cell suspensions from peritoneal lavage, spleens and bone marrow were then lysed for removal of red blood cells using ACK lysis buffer (Quality Biological, MD, 118-156-101). Cells were further resuspended in ice cold FACS buffer (2% FBS + 2mM EDTA) in 1x PBS and filtered through 40μm cell strainers to prepare single cell suspension. Immune subsets from colon mucosa were isolated according to previously published protocols ^63^. Briefly, dissected colons were opened and rinsed with 1x PBS to remove the luminal contents. To isolate intra-epithelial (IE) fraction, colon tissue was digested with 25 ml of fresh calcium/magnesium-free PBS containing 5 mM EDTA (Corning) and 2 mM DTT (Sigma) at 37°C with agitation for 20 min. IE fraction was removed by filtering through 100μm strainer. For lamina propria (LP) fraction, remaining colon tissue was digested with DMEM containing Collagenase IV (0.2 mg/ml), DNase I (0.05 mg/ml) supplemented with 10% FBS at 37°C for 60 min. The digested tissue was then gently mashed through a 100μm cell strainer. Cell pellets were resuspended in ice cold FACS buffer and filtered through 40μm cell strainers to prepare single cell suspension. For flow cytometry analysis on BMDMs, cells were harvested using ice cold detachment buffer containing 1XPBS, 2% FBS, and 2mM EDTA (Invitrogen, NY, 15575-038) and used for staining.

For staining surface markers, cell pellets were incubated with a fluorescent conjugated antibody cocktail for staining at 4°C for 30 min and were fixed using 1% paraformaldehyde (Sigma) for 20 min. For intracellular staining, cells were stimulated with PMA-20ng/ml, ionomycin-1μg/ml, brefeldin A-3μg/ml, and monensin-2μM and incubated at 37^0^C for 4 hours. After incubation cells were filtered through 40μm cell strainers. Cell pellets were incubated with antibody cocktail for surface markers at 4°C for 30 min and then fixed using True nuclear transcription factor buffer set (Biolegend, Cat no.424401) for 20 min at 4°C. Cell pellets were incubated with antibody cocktail for intracellular markers prepared in permeabilization buffer and incubated at 4°C for 30 min. All the antibodies were used at 1:400 dilution. Samples were then washed and resuspended in 1x PBS and acquired on BD FACS Symphony flow cytometer. Data were analyzed using FlowJo v10 (Treestar Inc.).

### Cholesterol quantitation

BMDMs (cultured and harvested as described earlier) were treated with HA (100uM) or kept untreated for 1hr and then stimulated with LPS plus IFNγ (100ng/ml) for 24hrs. Cell lysates were prepared using RIPA buffer (Cell signaling technology; Cat.9806S). Cholesterol estimation was performed using commercially available Amplex Red kit (Invitrogen, Cat. A12216) according to manufacturer’s instructions.

### RNAseq analysis

Total RNA was extracted from BMDMs using RNeasy Plus Mini Kit (Qiagen, 74136). RNA quality was validated using the TapeStation High Sensitivity RNA ScreenTape (Agilent). Libraries were prepared using Illumina Stranded mRNA Prep kit (500 ng DNase1 treated total RNA was used). Final library QC was done on the bioanalyzer using High Sensitivity DNA kit (Agilent). Next Generation Sequencing was performed with 100 bp paired end, on NovaSeq 6000 (Illumina) using SP v1.5 200 cycle kit. RNA-seq data were aligned using bowtie2 ^64^ algorithm against hg19 human or mm10 mouse genome version and RSEM v1.2.12 software ^65^ was used to estimate read counts and RPKM values using gene information from human Ensemble transcriptome version GRCh37.p13 or mouse Ensemble transcriptome version GRCm38.89. Raw counts were used to estimate significance of differential expression difference between any two experimental groups using DESeq2 ^66^. Overall gene expression changes were considered significant if passed FDR

<5% thresholds with additional fold change cutoffs when stated. Gene set enrichment analysis was done using QIAGEN’s Ingenuity® Pathway Analysis software (IPA®, QIAGEN Redwood City,www.qiagen.com/ingenuity) using “Canonical pathways”, “Diseases &Functions” and “Upstream Regulators” options. Selected results that passed the FDR <5% threshold were considered unless stated otherwise. Additional cutoffs on predicted activation state Z-score were applied where specified. Pathway clustering analysis was performed using Cytoscape ^67^. Selected pathways or functions network was imported with edges weight between any two pathways calculated as % of overlapped genes with overlaps of <20% not considered. Clustering was performed using Cytoscape plugin clusterMaker2 ^68^ with “Community Cluster (GLay)” option.

### Metabolomic analysis

Standards used for the metabolomic analysis include Hippuric Acid (Sigma-Aldrich, 112003; HA) and Hippuric acid-(phenyl-13C6, 99%) (Supelco, 93739; HA-13C6). Metabolomic analysis was performed as described previously with modifications ^69^. Polar metabolites were extracted from serum samples with 10-times the volume of ice-cold 80% methanol spiked with 1 μM HA-13C6 as an internal standard for HA in isotope dilution MS. Liquid chromatography-tandem mass spectrometry (LC-MS/MS) was performed on a Thermo Scientific Q Exactive HF-X mass spectrometer interfaced with a Vanquish Horizon UHPLC system. Gradient LC separation (0.2 ml/min, 26 min run time) used a ZIC-pHILIC column (150 × 2.1 mm, 5 µm particle size, EMD Millipore, 150460) at 45 °C. Mobile phase solvent A was 20 mM ammonium carbonate, 5 µM medronic acid, 0.1% ammonium hydroxide, pH 9.2, and solvent B was acetonitrile. All samples were analyzed by full MS with polarity switching. A quality control (QC) sample pool was also analyzed by data-dependent MS/MS with separate runs for positive and negative ion modes. Full MS scans were acquired at 120,000 resolution with a scan range of 50-750 m/z for positive ion mode and 65-975 m/z for negative ion mode. Data-dependent MS/MS scans were acquired for the top 10 highest intensity ions at 15,000 resolution with 1.0 m/z isolation width and stepped normalized collision energy (NCE) of 20/40/60.

Global metabolite profiling was performed using Compound Discoverer 3.3 SP1 (Thermo Fisher Scientific). Periodic technical injections of the QC pool were used to correct metabolite quantifications over the course of a batch analysis. Absolute quantitation of HA was performed using TraceFinder 4.1 (Thermo Fisher Scientific). Peak areas were determined for [M-H]-adducts in full MS scans with 10 parts per million (ppm) mass tolerance for HA (178.05096 m/z) and HA-13C6 (184.07112 m/z). Peaks were selected for integration based on retention times determined from the pure standards. Metabolite concentrations were determined from endogenous to internal standard ratios and calibration curves with linear fit and 1/X weighting. For both types of analyses, serum data were normalized per volume serum injected, which was equivalent for all samples.

### Lipidomic analysis

BMDMs were treated with hippuric acid and either LPS+IFNγ or IL-4. Media was removed from the plates by aspiration and cells were gently washed with chilled sterile PBS. Ice-cold methanol (OmniSolv, EMD Millipore, part # MMX04801) was used to quench and harvest the cells, which were subsequently transferred into glass tubes with PTFE-lined caps (Fisher Scientific 99502-10 and 999815). Samples were spiked with EquiSPLASH LIIDOMIX containing stable isotope-labeled internal standards representing 13 lipid classes (Avanti Polar Lipids, 330731). Lipids were extracted with chloroform/methanol/0.88% sodium chloride 2:1:1, and the aqueous layer was reextracted with synthetic organic phase, as described previously ^70^. LC-MS/MS was performed separately for positive and negative ion modes on a Thermo Scientific Q Exactive HF-X mass spectrometer interfaced with a Vanquish Horizon UHPLC system. Gradient LC separation (0.35 ml/min, 40 min run time) used an Accucore C30 Column (150 x 2.1 mm, 2.6 µm particle size, Thermo Scientific, 27826-152130) with 50:50 acetonitrile/water and 88:10:2 isopropanol/acetonitrile/water mobile phase solvents, both containing 5 mM ammonium formate and 0.1% formic acid. MS1 scans were acquired at 120,000 resolution, and data-dependent MS2 scans were acquired for the top 20 ions at 15,000 resolution with 0.4 m/z isolation width and stepped NCE of 20/30 for positive ion mode and 20/30/40 for negative ion mode. Lipid species were identified and quantified using LipidSearch 4.2 (Thermo Fisher Scientific). Identified lipid species were filtered by expected adduct and annotation grade based on lipid class, average peak quality across all samples, and variation of quantitation for triplicate injections of a QC sample pool. Quantification was based on MS peak areas, which were normalized to the internal standards for represented classes. Data were further normalized to the total lipid signal in each sample.

### Statistical analysis

Statistical analysis was performed using GraphPad Software version 10. To compare differences between two groups, two-group t-test or Wilcoxon rank sum test was used, and to assess differences between multiple groups, One-way ANOVA, Two-way ANOVA test with post-hoc multiple comparisons was used. Post-hoc multiple tests with Bonferroni’s procedure was applied to control type I error. Statistical analysis of non-targeted metabolomic and lipidomic data sets was performed using Perseus 2.0.9.0 ^71^. Data were log2-transformed, and pairwise comparisons between experimental conditions used two-tailed, unpaired Student’s t-test with Benjamini-Hochberg false discovery rate correction for multiple hypothesis testing (q-value).

## Author contributions

R.S.S conceived the study, designed experiments, and supervised the research. G.M., S.A.B., M.S., S.K.R., S.P.G., and T.M. conducted experiments, G.M. and R.S.S. analyzed the data and contributed to preparing figures. Y.Y. and A.K performed bioinformatic analysis and prepared figures, Q.L. helped in statistical analysis, A.R.G. assisted in conducting metabolomic experiments and analysis, N.Z. provided reagents and expertise. G.M. and R.S.S. prepared figures and wrote the manuscript. All authors contributed to revisions and provided critical comments.

## Data and materials availability

Materials and information are available upon requests to the corresponding author

