## Supplementary figures and images for "Aromatic Microbial Metabolite Hippuric Acid Potentiates Pro-Inflammatory Responses in Macrophages through TLR-MyD88 Signaling and Lipid Remodeling"

### Supplemental Figure 1

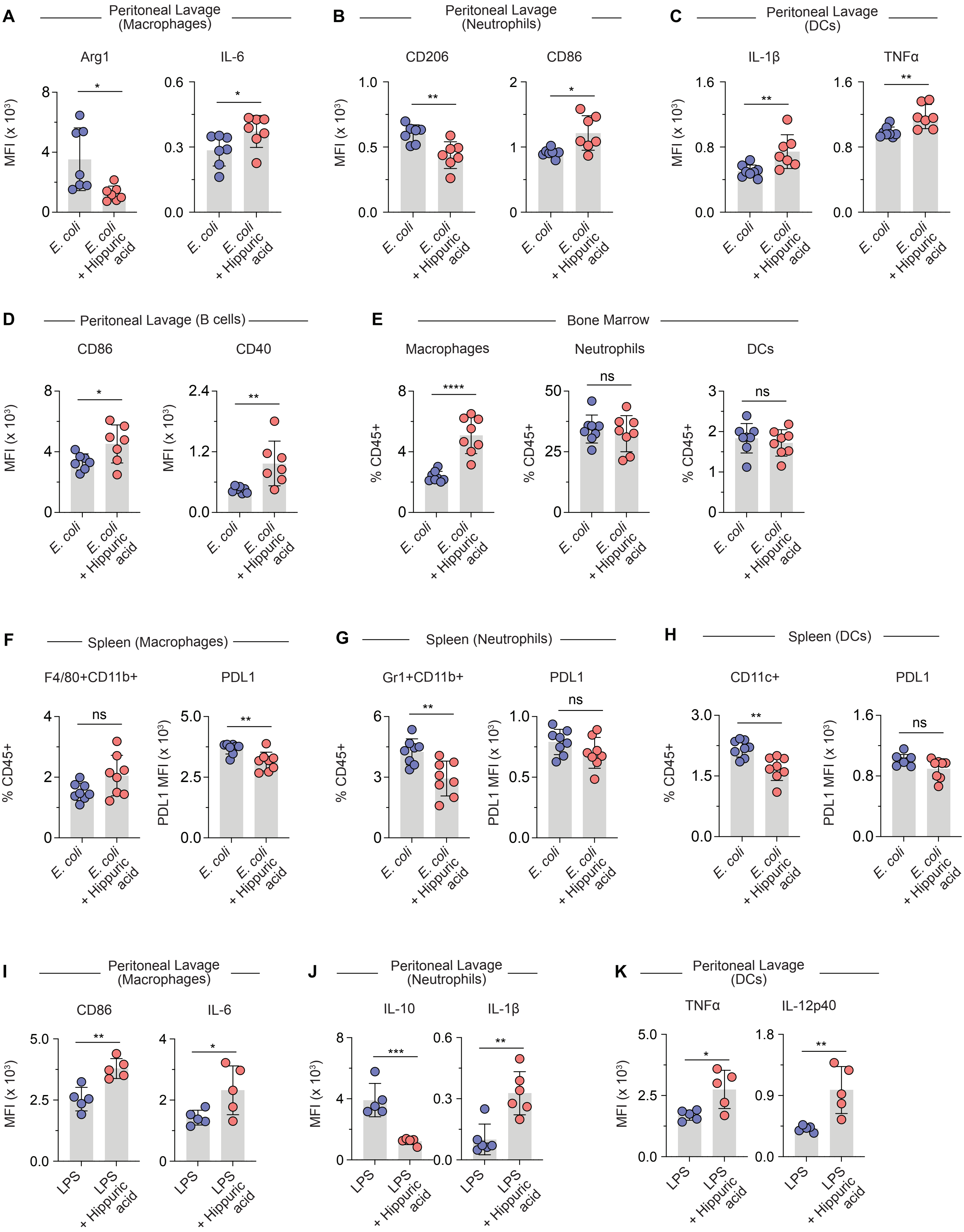

### Supplemental Figure 2

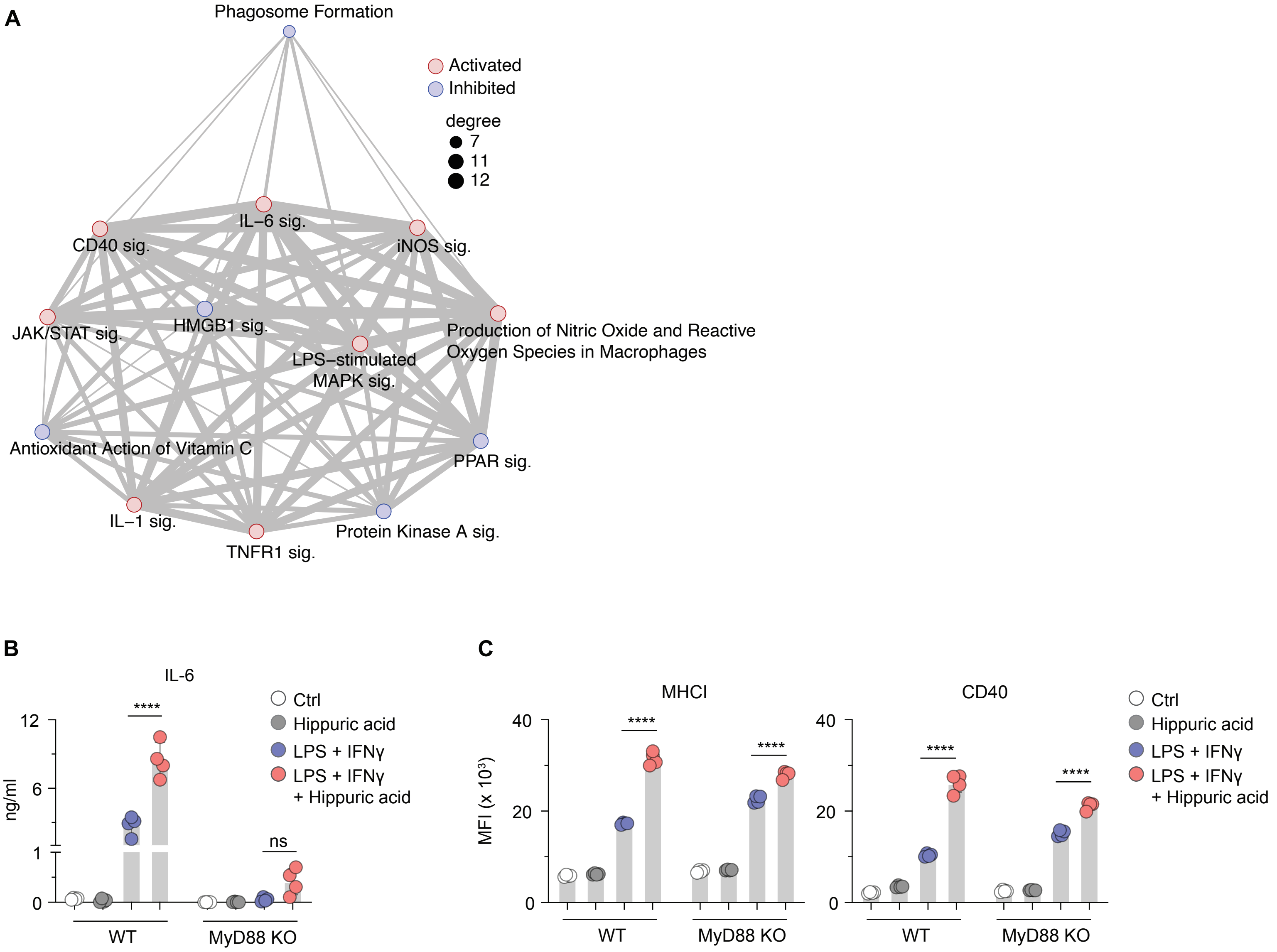

### Supplemental Figure 3

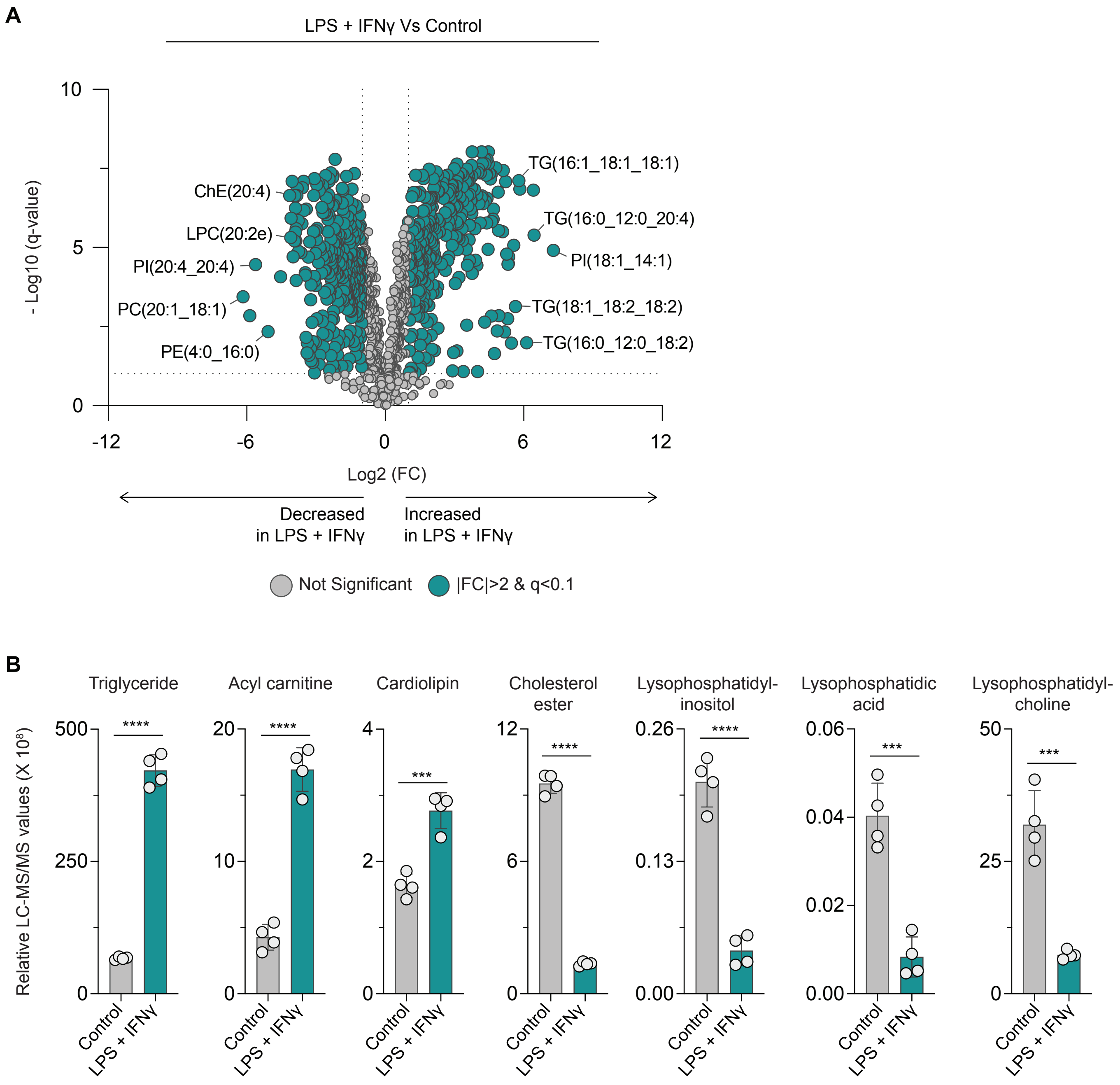

### Supplemental Figure 4

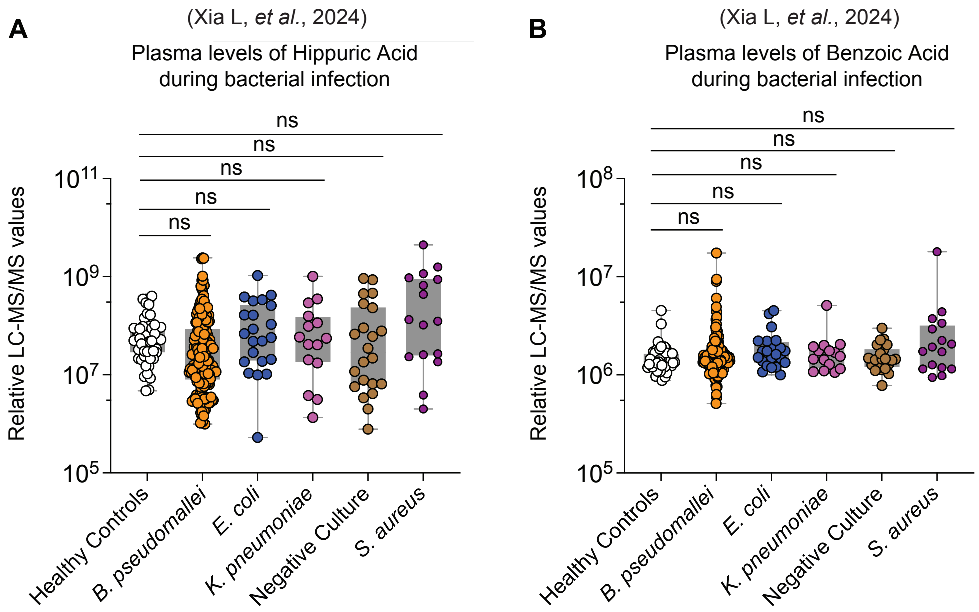
